# SOMDE: A scalable method for identifying spatially variable genes with self-organizing map

**DOI:** 10.1101/2020.12.10.419549

**Authors:** Minsheng Hao, Kui Hua, Xuegong Zhang

## Abstract

Recent developments of spatial transcriptomic sequencing technologies provide powerful tools for understanding cells in the physical context of tissue micro-environments. A fundamental task in spatial gene expression analysis is to identify genes with spatially variable expression patterns, or spatially variable genes (SVgenes). Several computational methods have been developed for this task. Their high computational complexity limited their scalability to the latest and future large-scale spatial expression data.

We present SOMDE, an efficient method for identifying SVgenes in large-scale spatial expression data. SOMDE uses self-organizing map (SOM) to cluster neighboring cells into nodes, and then uses a Gaussian Process to fit the node-level spatial gene expression to identify SVgenes. Experiments show that SOMDE is about 5-50 times faster than existing methods with comparable results. The adjustable resolution of SOMDE makes it the only method that can give results in ∼5 minutes in large datasets of more than 20,000 sequencing sites. SOMDE is available as a python package on PyPI at https://pypi.org/project/somde free for academic use.

## 1 Introduction

The spatial location of cells in tissues and corresponding gene expression profiles play pivotal roles in the study of tissue mechanism and tumor immune microenvironment. Spatial transcriptomics sequencing technologies provide gene expression profiles with spatial information, filling the gap between high throughput transcriptomics and their spatial context.

Since the publication of Spatial Transcriptomic (ST) (Ståhl *et al*., 2016) to today’s spatial transcriptomic sequencing technologies, the number of measured spatial data sites in one sample has increased from a few hundred to tens of thousands. Targeted in-situ sequencing technologies such as seqFISH+ (Eng *et al*., 2019) and MERFISH (Moffitt *et al*., 2018) capture single-cell level transcripts under the camera field of view (FOV). They splice multiple adjacent FOV results to form datasets with a large number of cells. Advanced Spatial Transcriptomic methods Slide-seq (Rodriques *et al*., 2019) and 10X Visium (the commercial version of original ST) can sequence 5,000∼25,000 spatial data sites with a resolution of 10 or 55 *μm*, respectively. It is foreseeable that the scale of spatial transcriptomic data will increase quickly.

Spatial transcriptomic data provides gene expressions with physical location information. Spatial variations of gene expression reflect the cell-cell interaction relationship (Dries *et al*., 2019) and help determine compositions of cell types that perform spatial specific functions. Identifying spatially variable genes (SVgenes) is the basic task for spatial transcriptomic data analysis. Compared with previous tasks of identifying highly variable genes (HVG) from gene expression profiles, SVgene identification needs to consider not only variations between cells but also the spatial significance of gene expression.

Several methods such as Trendsceek (Edsgärd *et al*., 2018), SpatialDE (Svensson *et al*., 2018), SPARK (Sun *et al*., 2020), scGCO (Zhang *et al*., 2018), Giotto (Dries *et al*., 2019) have been proposed for identifying SVgenes in recent years. Despite the great success of those methods in low-throughput spatial transcriptomic data, the high computational complexity hinders their application in large-scale datasets. For example, when the number of sites exceeds 20,000, Trendsceek, SpatialDE and SPARK will require at least 1000 minutes to get results (Sun *et al*., 2020).

To better cope with the growth of spatial transcriptomic data sizes, we present SOMDE, a scalable method to identify SVgenes with high computational efficiency. SOMDE uses the self-organizing map (SOM) neural network and Gaussian process to model spatial data. We conducted a series of experiments on both synthetic and real datasets which are obtained by two scalable protocols. Results showed that SOMDE gave similar SVgenes as existing methods, but in much less time with enhanced spatial pattern visualization. The adjustable number of nodes enables SOMDE to get results in large-scale datasets with more than 20,000 sites in only 5 minutes. We have developed SOMDE as a python package available on PyPI at https://pypi.org/project/somde free for academic use.

## 2 Methods

### 2.1 Overview of SOMDE

The transcript expression of SVgene is highly correlated with the spa-tial location and exhibits clustered, periodic or other patterns in space. Most of the published methods (Trendsceek, SpatialDE and SPARK) characterize the gene expression patterns based on the statistical spatial correlation models. The size of covariance matrices in their model grows quadratically as the number of spatial sites increases, and there-fore both time and memory consumption are needed to be optimized. Other methods (scGCO and Giotto) use gene expression binarization and non-statistical modeling to improve computational efficiency, but their scalability is still limited on the large-scale data.

The key idea of SOMDE is to construct a condensed representation of spatial transcriptomic data that both preserves the information of SVgenes and reduces the downstream computational complexity. We use the Self-Organizing Map (SOM) neural network to adaptively integrate neighboring data into different nodes, and then identify SVgenes based on the node-level spatial location and gene expression information using a modified Gaussian process (Fig.1).

**Fig. 1.**
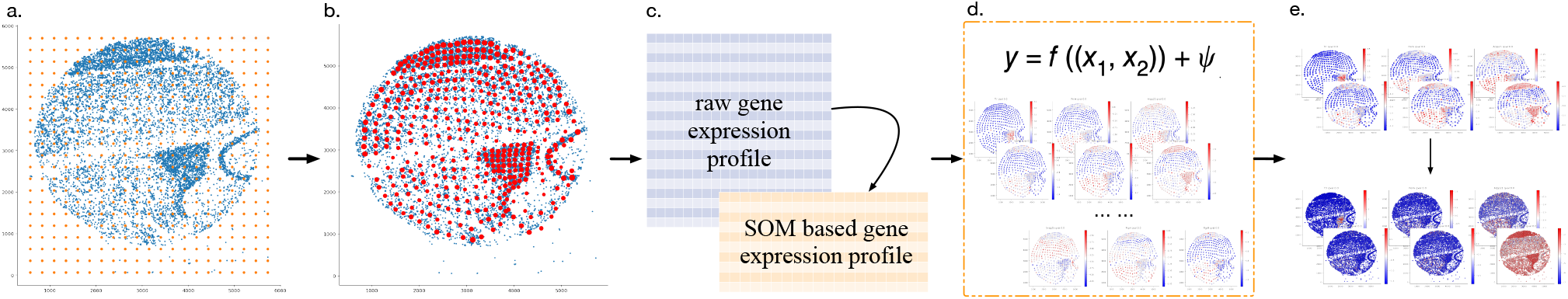
A schematic overview of SOMDE. (a) SOMDE initializes a self-organizing map (SOM) on the tissue spatial domain. (b) By training the SOM with data site locations, SOMDE merges the original data sites (blue spots) into different nodes (red spots). (c) Then SOMDE converts the original spatial gene expression to node-level gene meta-expression profiles. (d) SOMDE models the condensed representation of the original spatial transcriptome data with a modified Gaussian process. The variability of the gene expression *y* is decomposed into two parts, a spatial part f(x_1_, x_2_) that can be explained by the location of cells, and a non-spatial part Ψ for the rest of variability. (e) SOMDE identifies genes with high spatial variability expression patterns as SVgenes, and maps node-level expression patterns of SVgenes to their original expression patterns.

#### 2.1.1 Data site integration

The spatial expression of SVgenes usually has certain continuity. A proper strategy of data site integration should preserve the spatial expression patterns of SVgenes and the topological structure of data sites. SOM meets both the requirements. SOMDE first adopts SOM to integrate spatial data sites into nodes and then assigns the gene expression profile of each node.

SOM is an unsupervised neural network first proposed by Kohonen (Kohonen, 1984). It is an N×N array of neurons (nodes) arranged in a grid on the 2D plane. The neurons are connected to the input vector with trainable weights. The weights of neighboring nodes undergo competitive learning during the training phase so that a properly trained SOM will show the “self-organizing” property that preserves the topological relations and relative densities of the samples in the original input space (Kohonen, 1984; Zhang & Li, 1993). We apply SOM on the spatial gene expression data by using the 2D coordinates ***x*** of each data site as the input to the nodes. In this way, the trained SOM forms a condensed map that uses the node weights as a down-sampling representation of the original spatial information (Uriarte et al. 2005).

Specifically, we use a square SOM with *N* rows and *N* columns of nodes to learn the condensed representation. The size *N* can be determined according to the following formula:

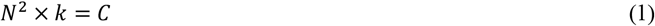

where *C* is a constant representing the total number of data sites, and k is the model parameter called neighbor number, which denotes the expected average number of original data sites each SOM node represents. This parameter controls how condensed the learned SOM is. A larger *k* means higher condensation.

We initialize the value and dimension of the SOM node weights ***M*** = (***m***_1_, ***m***_2_, …, ***m***_*N*×*N*_) with the uniform grid coordinates on the tissue spatial domain (Supplementary Materials). An example of the initialized SOM node weight is plotted in orange in Fig. 1(a). After initialization, SOM adjusts the weight of each node toward the tissue spatial topology through a repetitive training process using all spatial coordinates ***X*** = (***x***_1_, ***x***_2_, …, ***x***_*c*_) as training samples. We applied the batch SOM training algorithm (Wittek *et al*., 2017) in SOMDE. One spatial coordinate ***x*** is fed to SOM in one step and a cycle of all spatial coordinates is called one epoch. All node weights are parallelly updated once after one epoch. Multiple epochs are conducted during the whole training process. It takes less than 1 second to train a SOM in a dataset of 20,000 sites by taking this parallel implementation. The further elaboration of the training algorithm is in the Supplementary Material.

After training, each data site maps to a unique SOM node. Each SOM node represents the group of sites that mapped to it. Node weights in the learned SOM can be treated as new spatial coordinates for the data sites. The new spatial locations 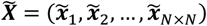 compose the sparse topology of the original data, as shown in red in Fig. 1b. To represent the expression of a gene on the condensed map, we define the gene “meta-expression” at a SOM node as the linear combination of the max value and average value of the gene expression in the group of sites that the node represents. The reason for not using average value as meta-expression is due to the high sparsity in large-scale data that could bias the result. Suppose *x*_*S*1_, *x*_*S*2_, … are a set of neighboring data sites mapped to a SOM node (*i, j*), the meta-expression 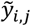 of one gene at this node is

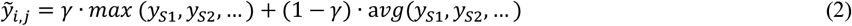

where *y*_*Si*_is the expression value of the gene at the data site *x*_*Si*_. The combination ratio *γ* balances the local maximum and mean gene expression. We use *γ* = 0.5 as the default in current experiments.

With this mapping and data integration for all gene and all data sites, we obtain the condensed representation of the original spatial transcriptome data by the meta-expression map in the SOM plane that best preserves the original topological and expression information. The resolution of this condensed spatial transcriptomic map can be adjusted by the parameter *k*.

#### 2.1.2 Spatial expression variability identification

The second step is identifying gene spatial expression variability with the condensed spatial transcriptomic map. We consider one gene meta-expression at one time and introduce an adjusted Gaussian process *H*_*G*_ in SOMDE to model the spatial correlation. Model *H*_*G*_ decomposes the expression variability into spatial and non-spatial components:

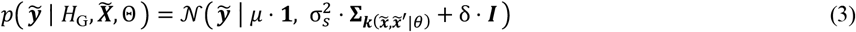

where 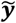 denotes one gene meta-expression in the SOM plane and 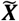 denotes SOM nodes locations. *δ*. ***I*** indicates the non-spatial variance given by Gaussian distributed noise in all observed gene meta-expression. 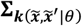 captures the spatial variation by using a selected kernel function ***k***(.). Here 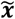 and 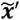 represent the locations of any two SOM nodes. The kernel function ***k***(.) introduces the spatial correlation between them. We applied a squared exponential (Gaussian) kernel in our model, since it can capture any spatial pattern (Svensson et al., 2018). By taking SOM nodes as the basic units, the covariance matrix is reduced to 1/*k*^2^ of the original matrix compared with previous statistical methods. We generated 10 Gaussian models with different length scales in SOMDE and choose the one with the highest log-likelihood ratio value. We used Θ to denote the parameters that need to be optimized, including the mean value *μ*, spatial signal variance 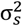 and noise variance δ. We applied gradient optimization to estimate these parameters.

The two separated variances allow us to get the fraction of spatial variation (FSV) in total variance by calculating the proportion of 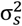 in 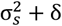 Compared with the original Gaussian process model, model *H* _*G*_ assumes one gene meta-expression at all spatial locations has the same mean value *μ* so that the non-spatial variance cannot be regressed out. Thus the maximum likelihood value of *H*_*G*_ mainly depends on spatial variance.

We use a similar log-ratio test as SpatialDE to determine the statistical significance of each gene’s spatial expression variability. Consider a Gaussian model *H*_0_ without the spatial covariance term:

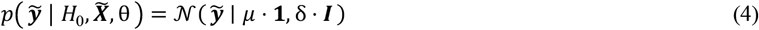

where *μ* and *δ* represent the mean value and noise variance, respectively. The log-ratio test between *H*_0_ and *H*_*G*_ reflects the significance of the spatial variance. And the P-value can be estimated by assuming the log-likelihood ratios (LLR) are χ2 distributed with one degree of freedom. SOMDE takes the LLR value as the spatial variability score for each gene and ranks them accordingly from high to low.

The top SVgenes ranks given by SOMDE at different resolutions may be different. To get a unified result, we propose a *k*-free solution called combined-SOMDE (cSOMDE) to merge all results. For each gene, cSOMDE takes the geometric mean of ranking results given by SOMDE with three different *k* as the final rank. This strategy considers the spatial variability of the same gene expression at different reduced resolutions.

SOMDE is implemented in Python and uses somoclu V1.7.5 for multi-thread building SOM. We choose SpatialDE V1.1.0, scGCO V1.1.1, SPARK V1.1.0 and Giotto V0.3.1 as representatives of statistical methods and non-statistical methods for comparison. All experiments are based on 16 AMD Ryzen 7 1700X Eight-Core Processors.

## 3 Results

### 3.1 Datasets and Experiments Design

We conducted a series of experiments on simulated data and real data to study the performance of SOMDE. The simulation data were used to evaluate the statistical power under different settings of signal-to-noise ratios (SNRs) and dropout rates. On the real data where ground truth is not available, we studied the numbers of SVgenes and examples of the identified SVgenes in comparison with other methods to illustrate the performance of the SOMDE.

#### 3.1.1 Experiments on Simulation Data

We adopted the Gaussian process (equation 3) to synthesize the simulation data. We generated a series of simulation data and used the 9,650 spatial locations (data sites) collected from the Slide-seq near Hippocampus (nHipp) (Rodriques *et al*., 2019). We generated 100 SVgenes and 900 genes with no spatial variation for all simulations. The whole generation procedure is similar to the SPARK (Sun *et al*., 2020), with details provided in the Supplementary Materials.

To mimic the real gene spatial expression data, we considered two data features, the signal-to-noise ratio (SNR) and the dropout rate. The SNR is controlled by the spatial (signal or pattern) and non-spatial (noise) variance in the Gaussian process model, and dropout is added by applying binary sampling of the data generated by the model.

We conducted three sets of experiments with three synthetic data of each. The first set of experiments aims to examine the performance of the methods under different SNRs when all information is observed, i.e. without dropout; To evaluate the performance when different combinations of SNRs and dropout rates exist in the data, we introduced the same dropout rate to data with different SNRs in the second set of experiments, and changed the dropout rate in data with the same SNR in the last set of experiments.

Specifically, in the first set, the combination of variance (signal variance, noise variance) we used are (0.1, 0.1), (0.1, 0.35) and (0.15, 0.15). The variance mentioned here is modeled on the log-normalized gene expression. The second set of experiments used the same settings of variance as the first set, but added a dropout rate of 0.2 to each simulation. In the third set of experiments, we fixed the signal and noise variances as 0.15 and 0.1, respectively, and used dropout rate of 0.2, 0.7 and 0.9 in the three simulations.

We compared SOMDE with three existing methods, SPARK, SpatialDE and scGCO. Giotto (SilhouetteRank) was not included because it provides only rank of genes without a specific cutoff for selecting the expected SVgenes. On each simulation data, we drew the FDR-Power curve to evaluate the performance of different methods.

#### 3.1.2 Experiments on the Real Data

We tested SOMDE on six large-scale real datasets: 10X Visium Brain, Visium Breast Cancer, Hippocampus (Hipp), near Hippocampus (nHipp), Liver and Kidney. These datasets are produced by two scalable spatial sequencing protocols. The Brain and Breast Cancer datasets are obtained by 10X Visium, and the other four datasets are obtained by Slide-seq (Rodriques *et al*., 2019). Brief descriptions of these datasets are shown in Table 1. The spatial resolutions of the 10X Visium and Slide-seq protocol are 55*μm* and 10*μm*, respectively. Although these data are not at the single-cell level, the measured gene expressions are highly correlated with spatial locations. The large gene and data site number are suitable for verifying the computational efficiency of our SVgenes identification method.

**Table 1.** The overview of datasets used in our experiments. Six datasets sequenced by two different protocols are used in our experiments. 10X data is downloaded from official 10X Genomics websites. The gene number and data site number indicate the scale of data after quality control.

| Protocol | Tissue | Gene number | Site number |
| --- | --- | --- | --- |
| 10X Visium | Brain | 14,414 | 2,698 |
| 10X Visium | Breast Cancer | 16,317 | 3,987 |
| Slide-seq | Hippocampus | 3,235 | 12,218 |
| Slide-seq | near Hippocampus | 2,555 | 9,650 |
| Slide-seq | Liver | 2,238 | 17,454 |
| Slide-seq | Kidney | 4,650 | 22,860 |

We applied different pre-processing strategies to these datasets. Af-ter getting the pre-filtered 10X Visium matrix, we filtered out genes whose total counts are less than 50 counts. Slide-seq datasets were downloaded from SpatialDB (Fan et al., 2019) website and were also pre-filtered. For Slide-seq data, we kept all the genes but removed the data sites where the total counts are greater than a 4-fold change of the average total counts. Then we log-transformed the raw count matrix and regressed out the total counts of each data site. The regressed matrix is the final normalized matrix for downstream analysis.

We set the default neighbor number *k* to 20 for all these datasets without deliberate selection. SOMDE takes spatial information and normalized gene expression matrix as input, gives the SVgene rank and tests the statistical significance of spatial variability with q-value. Genes with q-value smaller than 0.05 are identified as SVgenes. The SOMDE running time of all the above experiments has been recorded.

We further confirm the validity of identified SVgenes in nHipp and 10X Brain datasets from the biological and mathematical views. SOMDE and its variants cSOMDE were compared with the existing four methods. They are SpatialDE that uses the modified Gaussian process directly for identification, SPARK that uses the GLSM model, scGCO that uses graph-cut and Giotto (SilhouetteRank) that introduces the silhouette score to rank binarized expressed genes. For the cSOMDE method, we set neighbor numbers *k* =5, 20, 40 and *k* =4, 20, 40 in 10X Brain and nHipp datasets, respectively. The default settings were used for all other methods.

We compared the identified SVgenes numbers of SOMDE, scGCO, SPARK and SpatialDE. Besides the SVgene number, the rank similarities of different results are also important for validating the performance of methods. We compared the similarity of the top-ranked genes in the identification results of all methods, visualized the SVgene expression patterns and used ISH datasets for further verification.

### 3.2 Simulation results

In the first set of experiments (Fig. S1), we found that without dropout SPARK and SpatialDE both achieved extremely high power. SOMDE gained 0.8 power only with false discovery rate smaller than 0.05, regardless of the noise level being 0.1 or 0.35. Compared with the scGCO method which also focuses on SVgene identification on the large-scale data, SOMDE has much higher power. SOMDE can get the same power as the SPARK and SpatialDE after slightly increasing the signal variance.

We added the dropout rate of 0.2 in the second set of experiments (Fig. 2.). Under the same SNR as the experiment set 1, the power of all methods decreased (SPARK failed to give results on the dropout simulations due to its sensitivity to data sparsity). SOMDE gets the power higher than 0.9 after increasing the signal variance to 0.15, and has a better performance than scGCO in all simulations.

**Fig. 2.**
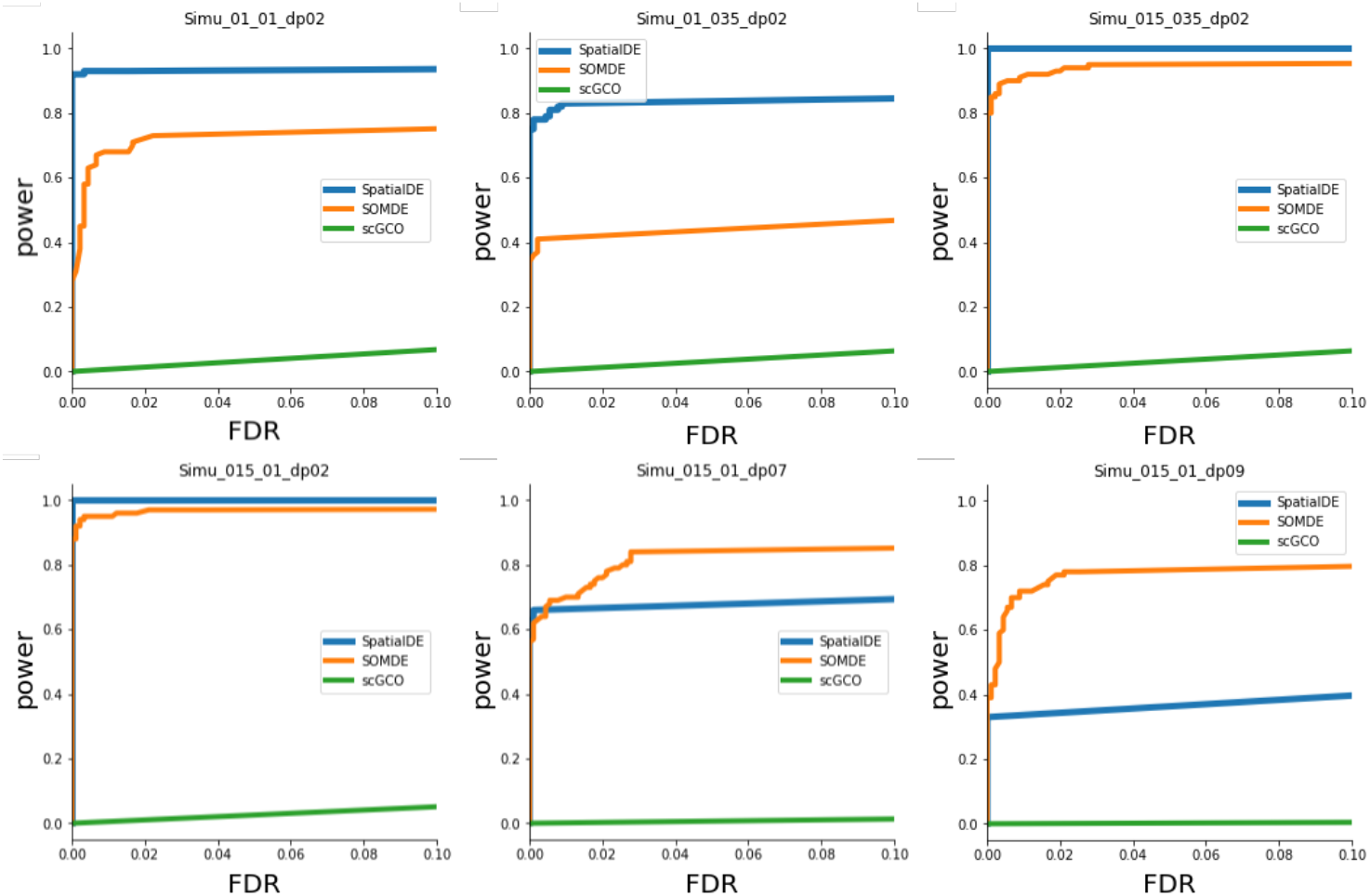
Performance of different methods in simulation data with different signal-to-noise ratios (SNRs) and dropout rates. To mimic the real cases, we adjusted the SNR and dropout rate in the simulation data. We used the FDR-Power curve to show the performance of different methods on each simulation. The first row of figures gives the results under different SNRs with a fixed dropout rate (experiment set 2). The second row shows the results when the dropout rate varies while SNR is fixed (experiment set 3). SNR is controlled by two parameters, the signal (pattern) variance and the noise variance. The specific parameter settings (signal variance, noise variance, dropout rate) of the six results are (0.1,0.1,0.2), (0.1,0.35,0.2), (0.15,0.35,0.2), (0.15,0.1,0.2), (0.15,0.1,0.7), (0.15,0.1,0.9).

In the third set of experiments, we found that SOMDE outperforms other methods in addressing data sparsity (Fig. 2.). SOMDE is more powerful than SpatialDE and scGCO across all FDR cutoffs in the simulation with the dropout rate being 0.9. The result shows that the data integration step in SOMDE not only boosts the computation but also enhances the signal of patterns on the large-scale sparse data.

SOMDE exceeded other methods in computational efficiency. On the simulation data with 9650 data sites and 1000 genes, SOMDE spent 20 seconds finishing all the calculations. As comparisons, it took 450 seconds for scGCO, 30 minutes for SpatialDE and more than 3 hours for SPARK to get results on the data.

### 3.3 Real data results

SOMDE gave the SVgene rank and the SVgenes number in each dataset (Fig. 3. & Supplementary Materials). In the 10X Brain and Breast Cancer datasets, SOMDE found 5,455 and 7,356 SVgenes, respectively. In the Liver dataset, the SVgenes number is the smallest with only 164 genes. SOMDE found 699, 379, 522 SVgenes in Hipp, nHipp and Kidney datasets, respectively. *CRISP3, CPB1, CXCL14, COX6C* and *FCGRT* are the top 5 SVgenes identified by SOMDE in the 10X Breast Cancer dataset. The top 5 SVgenes in the 10X Brain dataset are *Sparc, Agt, Slc6a11, Tcf7I2* and *Nrgn*, respectively. *Nrgn* and *Tcf7l2* were also identified as the top 10 SVgenes in the nHipp and Hipp dataset,. *Ttr* gene has the most spatial expression variability in both nHipp and Hipp datasets. In Liver and Kidney datasets, the top 5 SVgenes are *Mup17, Hpx, Mgst1, Glul, Mup3* and *Napsa, Kap, Aadat, Mpv17l, Acadm*, respectively. All SVgene rank lists of these six datasets are available in the Supplementary Material.

**Fig. 3.**
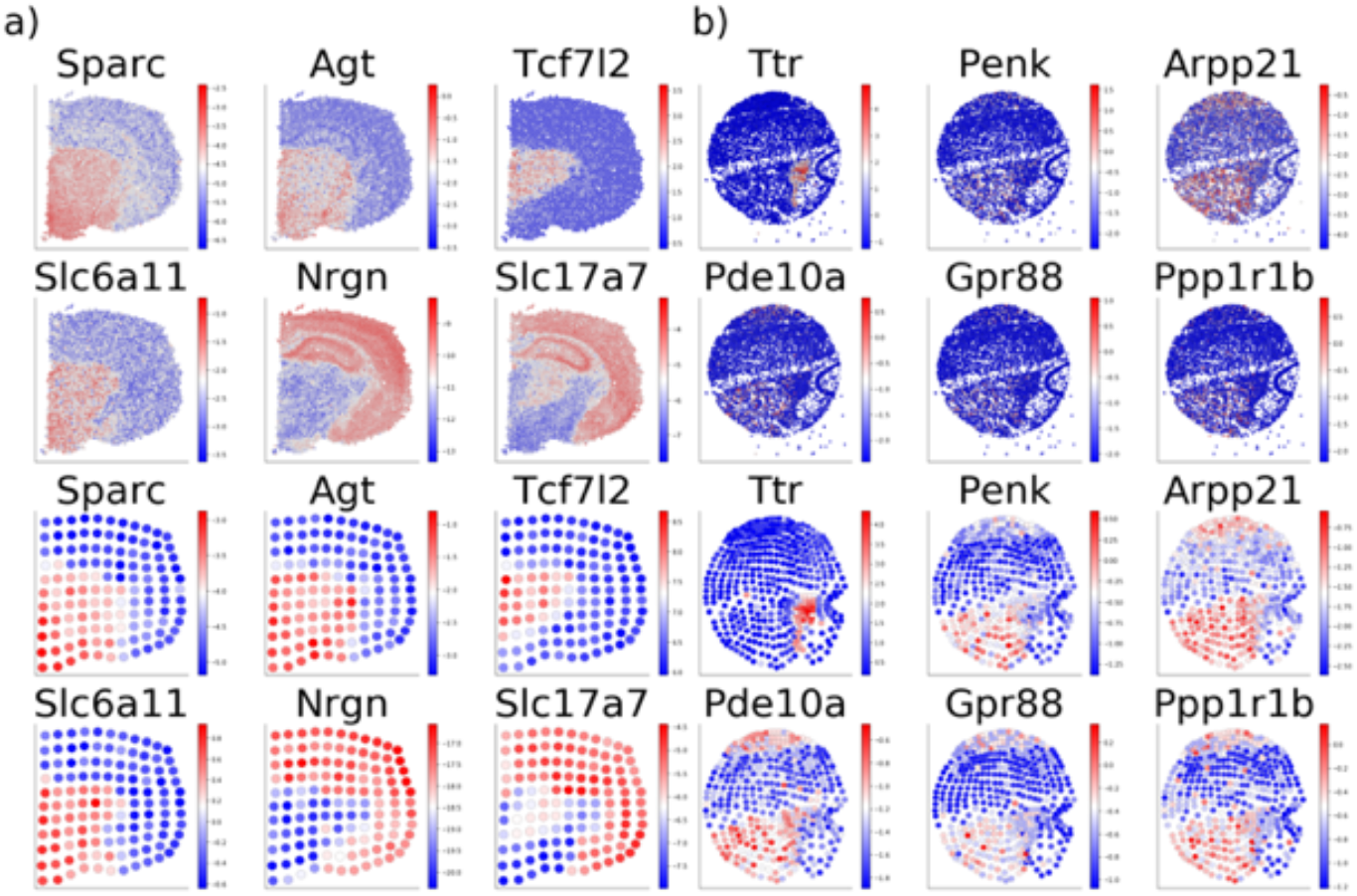
The top 6 SVgenes found by SOMDE are visualized in original and SOM space. Top 6 SVgenes found in 10X Brain and nHipp datasets. Each subplot denotes one gene spatial expression pattern. Color implies the relative expression value within the plot. The first two rows and the last two rows are in the original space and SOM space, respectively. (Zoom in for details

From the results, we found that the different proportions of SVgene among datasets are more likely due to the biological characteristics of the tissue, rather than technical bias or different data size. In tissues with complex spatial correlated functions such as the whole adult mouse brain or hippocampus, the proportion of SVgenes is far greater than in the adult mouse kidney or liver tissues.

We plotted the top 50 SVgenes spatial expression patterns at the original resolution and node-level resolution in all datasets. (Fig.3 & Supplementary Figures) Each spot in the figure denotes one data site or one SOM node, and the color represents the relative expression values within one plot. The visualization results on the original data and condensed map show the similar topological structure of data sites and spatial gene expression patterns. We also checked the bottom 50 SVgenes on 10X Brain data (Fig. S11). The spatial variabilities of these genes in the original resolution are less clearly compared to that of the top 50 ones and only a few are apparent to human eyes. However, most of these patterns got enhanced in the condensed map of SOM (Fig. S11).

The integration step of SOMDE improves the SNR of gene expressions on the Slide-seq nHipp dataset. *Pde10a, Gpr88* and *Ppp1r1b* genes (Fig. 3b) have low expression levels in the original tissue spatial domain so that it is hard to visualize their high spatial expression variability due to the noise and sparsity. On the condensed map, however, these genes show strong spatial patterns and variabilities. SOMDE successfully enhanced gene expressions via the combination of the maximum and average local expression. These results also suggest that the visible patterns shown on the original spatial domain may not be a golden standard for determining spatial variation.

We further analyze our SOMDE results on the Slide-seq nHipp and 10X Brain datasets to verify the identification results. We use the MGI database (Smith *et al*., 2019) and Allen Brain Atlas (Lein *et al*., 2007) as references to verify the genes’ morphological function and spatial expression distribution. SVgenes identified by SOMDE showed clear spatial variation and well consistent with existing In Situ Hybridization (ISH) patterns of the same tissue (Fig. S9 & 10). For example, *Nrgn*, one of the top 5 SVgenes in the 10X Brain dataset, is responsible for the transcription of neurogranin protein that encodes transcription and signal synapse transduction. Expression patterns on both 10X Brain and ISH data reveal that *Nrgn* mainly expressed in the cerebral cortex and hippocampal formation (Fig. S9). *Camk2n1*, another SVgene identified in the mouse brain, is responsible for the production of enzyme regulators and is highly expressed on the outer and posterior parts of the cerebral cortex on our condensed map. This spatial pattern is also cross-validated by the ISH data (Fig. S9). Both *Nrgn* and *Camk2n1* play important roles in synaptic long-term potentiation (Ling, K.H., 2011), and they have a similar spatial variability score and SVgene rank, suggesting our results are of potential biological interest.

We also confirm the validity of SOMDE from the mathematical view. Fig. 4 shows the log-likelihood ratio (LLR) and the fraction of spatial variation (FSV) of all genes given by SOMDE in 10X Brain and nHipp datasets. Each spot denotes one gene and we highlight the spots of the top 5 SVgenes. The top 5 SVgenes in both datasets identified by our method have high FSV and LLR values. *Sparc* gene has the maximum LLR value corresponding to the most significant spatial patterns shown in Fig.3. For highly spatially variable genes, LLR and FSV values are positively correlated as expected. The gene spot distribution indicates most spatially variable genes have significant statistical values, which shows that the integration of spatial sites does not affect the identification of these gene variabilities.

**Fig. 4.**
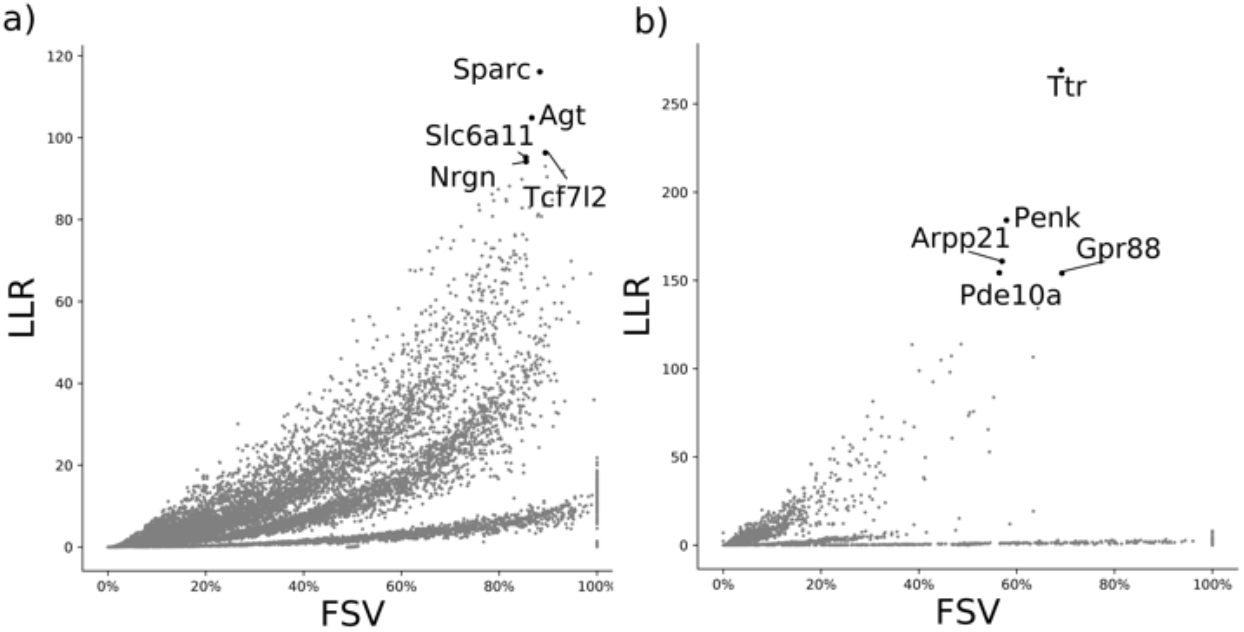
Quantitative results of SOMDE in 10X Brain and nHipp datasets. The x-axis represents the fraction of variance explained by spatial variation (FSV), and the y-axis represents the log-likelihood ratio (LLR). The top 5 genes selected by our method are high on both indicators. Subfigure a) and b) show results in brain datasets and nHipp datasets.

Table 2 shows the running time of SOMDE in all six datasets. Running time refers to the total time from training self-organizing map to obtaining SVgene ranks and spatial variability scores. The adjustable map size guarantees the condensed maps are at the same resolution on different datasets. SOMDE gave all the results with no more than 5 minutes regardless of the gene and data site number. We can conclude that the condensed transcriptomic maps make the computational efficiency of SOMDE not limited by the dataset size.

**Table 2.** the running time of SOMDE in multiple datasets. The results of the SOMDE on different datasets with *k* = 20 (*k* = 10 on the 10X Cancer data). Although the running time is affected by the gene and sequencing site number, SOMDE gives results in less than 5 minutes in multiple datasets (some datasets even exceed 20,000 data sites), which shows the superiority of our method in terms of running time.

| Dataset | Gene num. | Site num. | Map | Time (sec.) |
| --- | --- | --- | --- | --- |
| 10X Brain | 14,414 | 2,698 | 11×11 | 247 |
| 10X Cancer | 16,317 | 3987 | 19×19 | 354 |
| Hipp | 3,235 | 12,218 | 24×24 | 56 |
| nHipp | 2,555 | 9,650 | 21×21 | 57 |
| Liver | 2,238 | 17,454 | 29×29 | 58 |
| Kidney | 4,650 | 22,860 | 33×33 | 147 |

### 3.4 Comparison with existing methods on the real data

#### 3.4.1 Gene ranking similarity

All methods get the results on the 10X Brain and nHipp data, except that SPARK failed to run on the nHipp data because of the data sparsity. These methods revealed similar judgments on the most spatially variable genes, regardless of their own criteria or principles (Fig. 5. & Supplementary Figures). In the 10X Brain dataset, SOMDE shared 4,809, 5,501, 3,351 common SVgenes with SPARK, SpatialDE and scGCO (Fig. 5a), respectively. We also compared the similarity of all the results with different cutoffs. *Agt, Nrgn, Sparc, Camk2n1, Slc17a7, Slc6a11* and *Tcf7l2* are at the intersection of the top 10 SVgenes identified by SOMDE and other methods. Among the top 500 SOMDE SVgenes, cSOMDE, SpatialDE, Giotto, scGCO and SPARK identified 435, 274, 277, 259 and 39 identical SVgenes, respectively (Fig. S12). Compared with other methods, our method and SpatialDE use similar statistical models to infer spatial variability. Thus SOMDE has a high consistency with the results of SpatialDE.

**Fig. 5.**
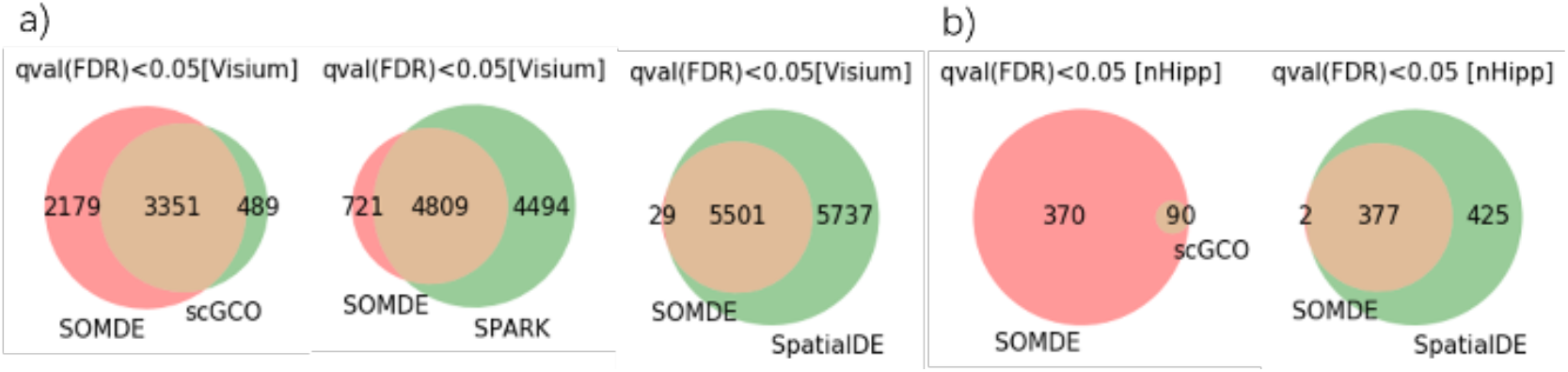
Number of SVgenes identified by four methods on the 10X brain (a) and nHipp data (b). Genes with FDR (q value) <0.05 are identified as SVgenes. Most of the 5530 genes found by SOMDE in 10X brain dataset were also identified by other methods. More than 99% SVgenes identified by SOMDE have also been identified by SpatialDE in the nHipp data. Since cSOMDE and Giotto cannot provide the SVgene number, we compared the top 500 SVgene of each method and Venn diagrams are in the supplementary figures.

We observed similar results in the nHipp data. SOMDE found all SVgenes identified by scGCO, and 377 SVgenes identified by SOMDE are shared with SpatialDE (Fig. 5b). Focusing on the rank similarity of the top SVgenes, we found that *Ttr, Mef2c, Pcp4, Pde10a, Penk, Arpp21* and *Enpp2* are the intersection genes among the top 10 genes identified by SOMDE and other methods. Among the top 500 SOMDE SVgenes, cSOMDE, SpatialDE, Giotto and scGCO identified 457, 401, 277, 142 and 93 identical SVgenes, respectively (Fig. S13). We concluded that SOMDE has a high overlap with other methods on the top SVgenes. Over half of SVgenes among the top 500 identified by SOMDE and other methods are identical, except scGCO only report 9 SVgenes in the nHipp data.

We further found that SOMDE and cSOMDE have almost identical SVgene ranks in both two datasets. They shared over 85% percent of common SVgenes cross any rank cutoffs (Fig. S12. &S13). The diagonal rank plot (Fig. S14) also shows that the cSOMDE result obtained from the geometric mean has a similar rank to that obtained from a single SOMDE result. Besides, we compared the SOMDE results under a set of adjacent map sizes on the 10X Brain and nHipp data (Fig. S15). The diagonal scatter points in the rank plots elaborated that the results with adjacent map sizes (parameter *k*) have high similarity. We also compared the SOMDE results under different parameters *γ*. Similar diagonal scatter points are shown in Fig. S18.

#### 3.4.2 Running time comparison

We selected three real representative datasets: 10X Brain, nHipp, and Kidney to test the computational efficiency of different methods on the large-scale data. The number of sequencing sites in these three datasets gradually increases, as shown in Table 2. The comparison results of the five methods are shown in Fig. 6a. SOMDE spent 247, 57 and 144 seconds to give results in the three datasets. It took scGCO 2,917, 400 and 2,424 seconds to get the results in these datasets, although it is designed to improve the computational efficiency. Giotto and SpatialDE spent more than 5,000 seconds on the Kidney dataset. SPARK spent more than 4 hours in the 10X Brain data and failed to give the result on the nHipp data so that we did not further test it on the Kidney data. The adjustable resolution of SOM makes SOMDE the only method that gets results in large-scale datasets in 300 seconds. Unlike other methods, the computational complexity of SOMDE barely increases with the growth of the dataset size, and is only affected by the number of genes. Therefore, the running time in Liver and Kidney datasets is even less than which on the 10X brain data.

**Fig. 6.**
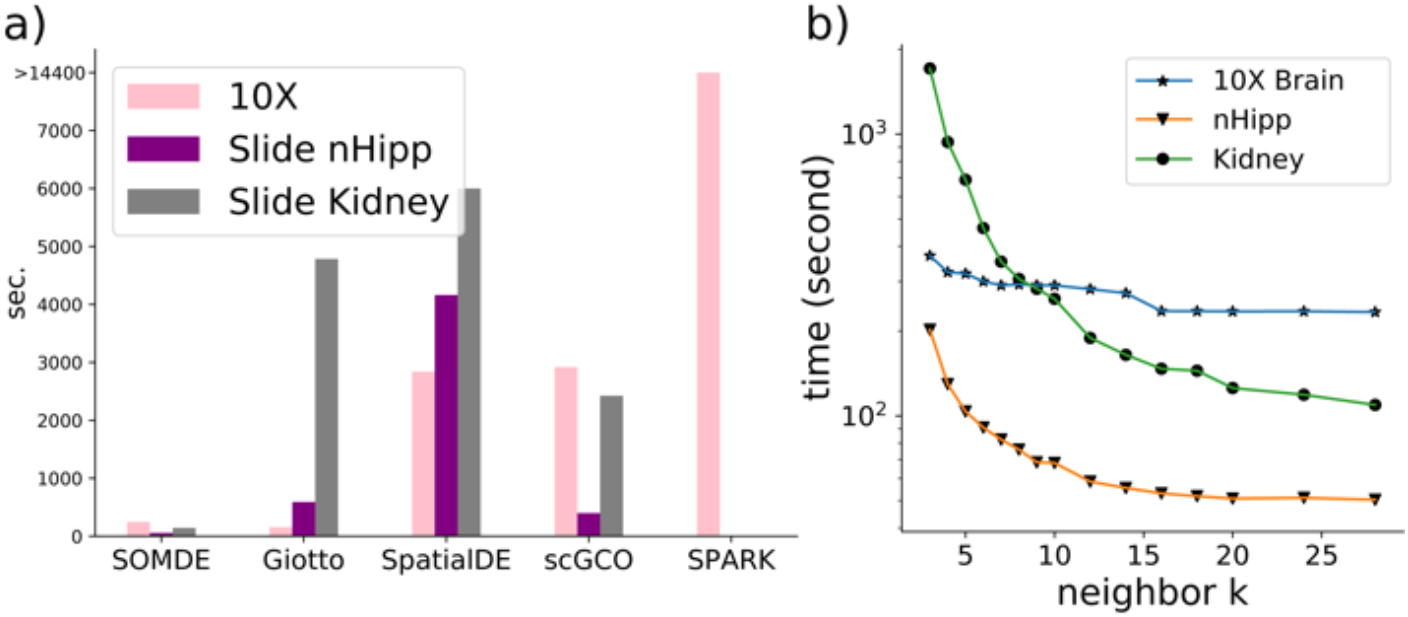
Running time Comparison. a) The total running time of 5 methods in 3 datasets with different sizes. Colors correspond to different datasets. b) Running time of SOMDE under different neighbor numbers. SOMDE gives results rapidly in all three datasets and has the most impressive improvements on the kidney dataset which has the largest number of spatial sites.

To explore how the parameter *k* affects the running time of SOMDE, we applied SOMDE with gradually changed k on the three selected datasets and recorded the running time (Fig.6b). From these results, we can see that a small decrease in the size of condensed map can lead to a significant improvement in the computation efficiency, as demonstrated by the sharp decrease at the beginning of the running time curves (Fig. 6b). It is also clear that the curve trends to stabilize as k gets larger enough. This is because for a large k, the condensed map becomes so small that modeling its covariance is no longer the main factor contributing to the whole running time of SOMDE.

Based on the above observations, we know that in real applications, a proper k should not be too small so that high computational efficiency can be guaranteed. Meanwhile, it should not be too large because a large k brings a small gain in the computational efficiency but a high risk of losing more local spatial patterns in the condensed map. We recommend *k*=5∼20 in SOMDE for most scenarios.

## 4 Discussion

The increasing throughput of spatial transcriptomic data enables researchers to study the gene expression in the spatial context at a larger scale, but brings computational challenges to bioinformatic tools as well. We present SOMDE, an SVgenes identification method that extends the application to the large-scale dataset by combining the advantages of machine learning and statistical models. SOMDE integrates gene expression and spatial locations to a condensed map by applying self-organizing map and identifies gene spatial expression variability via Gaussian Process.

Experiments on multiple datasets with different sequencing protocols showed that the SOM-based condensed map well preserved the topological structure and the gene spatial expression patterns. The SVgenes identified by SOMDE are highly consistent with other methods and existing gene ISH data. SOMDE also enhances the spatial patterns of SVgene expressions, which helps for better visualization. Experiments showed that choices on the condensed map size within a proper range do not change the identification results much. Therefore one SOMDE model can work for many scenarios.

It should be noted that the degree of spatial variation of a gene’s expression is a relative matter and defining the hard cutoff of SVgene and non-SVgene is not feasible. This is especially true when we consider different possible modes, scales and localization of the variation. The observed null distribution of test p-values for permuted data (Fig. S16 & S17) also confirms the intrinsic complexity of this problem. This is a topic that needs further research, which should be put into the context of downstream biological analyses of the spatial variations. SOMDE provides a relative ranking of SVgenes with regard to the strength and significance of the variation.

Compared with existing methods, SOMDE is more similar to those statistical models. This is related to our choice of the Gaussian Process as the significance test model. The self-organizing map can be considered as the frontend and Gaussian process as the backend of SOMDE. The backend statistical model can take any other forms such as the marked point process used in Trendsceek or GLSM used in SPARK. SOMDE can also be extended to 3D spatial sequencing data in the future, since the Gaussian Process and SOM have no restriction on dimensionality.

A key challenge in identifying SVgenes in large-scale spatial transcriptomics data is the high computational complexity. SOMDE tackles this challenge by building a condensed map from the original data. We should notice that this also introduces limitations on the application of the method. Although SOM provides adaptive down sampling that can better preserve the topology and densities in the original data than uniform down sampling, some local details can still be missed (Fig. S19). We may adopt a hierarchical strategy to first detect variations at the global view first and then at scan for local variations in smaller areas of interest to better capture SVgenes at multiple resolutions.

## Acknowledgements

We thank Prof. Guocheng Yuan, Dr. Qian Zhu and Dr. Ruben Dries for their helpful discussions. We thank the anonymous reviewers for their constructive suggestions.

