## Supplemental Material for "SOMDE: A scalable method for identifying spatially variable genes with self-organizing map"

#### Self-Organizing Map Initialization & Training Algorithm

After an $N \times N$ SOM is established, the SOM training algorithm takes the input data (spatial coordinates of data sites) $\boldsymbol{X} = (\boldsymbol{x}_1, \boldsymbol{x}_2, \ldots, \boldsymbol{x}_C)$ to iteratively update the node weights $M = (\boldsymbol{m}_1, \boldsymbol{m}_2, \ldots, \boldsymbol{m}_{N\times N})$.

##### Node weight initialization

The dimension of the SOM node weights is the same as that of the data sites' spatial coordinates. We initialize the SOM node weights with the uniform grid coordinates on the tissue spatial domain. Let $\boldsymbol{x}_i = (p_x^i, p_y^i)$, where $p_x^i$ and $p_y^i$ are the coordinates among the x and y axis, respectively. The extreme values of the coordinates are defined as follows:

$$x_{min} = \min(p_x^1, p_x^2, \ldots, p_x^n)$$

$$x_{max} = \max(p_x^1, p_x^2, \ldots, p_x^n)$$

$$y_{min} = \min(p_y^1, p_y^2, \ldots, p_y^n)$$

$$y_{max} = \max(p_y^1, p_y^2, \ldots, p_y^n)$$

Then the weight vector $\boldsymbol{m}_{i\times j}$ of node $n_{i,j}$ in the two-dimensional grid is defined as:

$$\boldsymbol{m}_{i\times j} = (x_{min} + \frac{x_{max} - x_{min}}{N} \times i, y_{min} + \frac{y_{max} - y_{min}}{N} \times j)$$

##### Training algorithm

The basic idea of SOM is training the node weight though the competitive learning. When a spatial coordinate is fed into SOM, the distance between it and the weight of each node will be calculated. The node whose weight vector is most similar to the input 'wins' the competation and is called the best matching node. The weight of this node and the nodes adjacent to it in the SOM grid will be adjusted to the input vector. And finally a properly trained SOM will show the "self-organizing" property that perserves the topological relations and relative densities of the samples in the original input space.

A graphic illustration is shown here: The upper layer contains SOM node with initialized weight, and the lower layer is the input data, which contains the coordinates of all data sites in the original tissue spatial domain. The node weights are adapted toward the tissue spatial topology through repeating training on the spatial coordinate $\boldsymbol{X} = (\boldsymbol{x}_1, \ \boldsymbol{x}_2, \ldots, \boldsymbol{x}_C)$. We want to mentioned that the node positions in the upper layer are plotted by there weights in this figure. While in other SOM figures, nodes positions only refer to the relative positions on the SOM grid and are unchanged. After proper training, Node weights form a condensed map of the original input data.

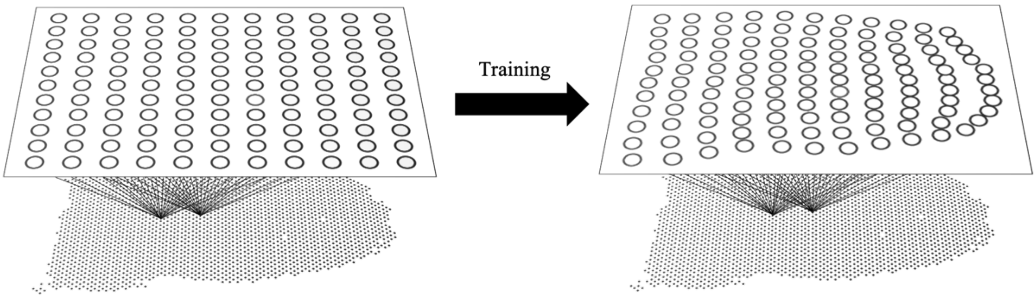


At a given step $t$, one spatial coordinate $\boldsymbol{x}(t)$ is selected to map to its best matching node $n_b$:

$$bm(\boldsymbol{x}(t)) = n_b \in M$$

such that:

$$d(\boldsymbol{x}(t), \boldsymbol{m}_b(t)) \leq d\left(\boldsymbol{x}(t), \boldsymbol{m}_j(t)\right) \ \forall \boldsymbol{m}_j(t) \in M$$

where $d$ is the distance function on the dataset in the tissue spatial domain. Then the node $n_b$ and its neighbors' weight vectors are adjusted toward the input data. In the traditional training algorithm, the adjustment follows this formula:

$$\boldsymbol{m}_j(t+1) = \boldsymbol{m}_j(t) + \alpha h_{b,j}(t)(\boldsymbol{x}_t - \boldsymbol{m}_j(t))$$

where $\alpha$ controls the learning rate, $h_{b,j}(t)$ is the neighborhood function that decreases for nodes further away from the best match nodes in the grid. Gaussian kernel is the most used neighborhood function:

$$h_{b,j}(t) = \exp\left(\frac{-||r_b - r_j||}{\delta(t)}\right)$$

where $r_b$ and $r_j$ represents for the grid coordinates of the respective nodes. $\delta(t)$ controls the area of influence and decreases from iteration to iteration.

This training procedure is repeated on the same dataset to increase the fit. Eventually, the node weights capture patterns in the input data and the map preserves the topological constraints and density relations among the input data in the original space.

This traditional training algorithm demands to set the initial value and attenuation function of the learning rate, which causes the instability of the training results. And it is also time-consuming. We applied the batch SOM training algorithm (Wittek *et al.*, 2017) in SOMDE. The updating formula for weight vector $m_i$ of node $i$ is:

$$m_i(t_C) = \frac{\sum_{t'=t_0}^{t'=t_C} h_{b,i}(t')\boldsymbol{x}(t')}{\sum_{t'=t_0}^{t'=t_C} h_{b,i}(t')}$$

where $m_i(\cdot)$, $h_{b,i}(t')$ and $\boldsymbol{x}(t')$ follow the same definitions in the above. The beginning and the end step of one epoch are represented as $t_0$ and $t_C$, respectively. One epoch contains $C$ (data site numbers) total steps and one spatial coordinate $x_i$ is fed to SOM in one step. We set the total epoch number to 10 following the default setting in somoclu (Wittek et al., 2017). This batch training form avoids the need for setting the hyperparameters, and node weights are parallelly updated by all input data after one epoch. Since each update of one SOM node only affects its neighboring nodes, the training process has lower computational complexity. We set the total epoch number to 10 following the default setting in somoclu (Wittek et al., 2017).

#### Simulation Generation

We first ran SOMDE on the original Slide-seq nHipp expression data to identify SVgene patterns. Genes with q-value smaller than 0.05 are filtered as SVgenes. Then we performed the automatic expression histology (AEH) function implemented by SpatialDE on all SVgenes. AEH grouped SVgenes with similar patterns into 10 clusters and generated the posterior expected expression values $\tilde{\boldsymbol{\mu}}$ for each cluster. Here $\tilde{\boldsymbol{\mu}}$ is the log-transformed expression value on the condensed map. To achieve the original resolution expression patterns, we assign the node expression values to the data site groups mapped to it. The 10 patterns identified by SpatialDE are shown as follows:

(based on the condensed map):

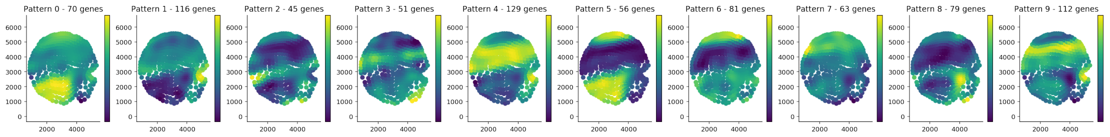


(based on the data sites):

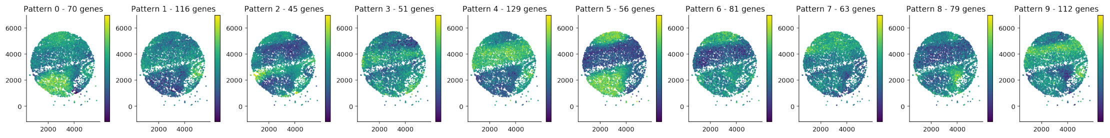


We scale the mean and variance of each expression pattern to -1.34 and 0.15, respectively. -1.34 is the mean values on the real log-transformed gene expression matrix. Then for each simulation experiment, we simulated 100 SVgenes samples by adding the Gaussian noise on these 10 patterns. For non-SVgenes, we directly sample the gene expression from the Gaussian distribution with the mean being -1.34 and variance being the same as the positive samples. After getting the total log-transformed 1000 gene expression on the 9650 data sites, regardless of whether it is SV or non-SV genes, we exponentiated the expression matrix $A \in \mathbb{R}^{1000\times9650}$ and simulated the gene expression count data based on a Poisson distribution $Poi(\lambda)$. The parameter $\lambda$ is a product of the exponentiated values in the matrix and the total read counts $N_j$ that is obtained from the read data. In other words, for gene $i$ at data site $j$, the expression count is sampled from $Poi(A_{i,j}N_j)$.

We added the dropout rate to simulate the sparseness on the Slide-seq data. Specifically, We randomly discarded the expression value of each gene on some nodes according to the dropout rate.

### Tables & Figures

**Table S1.** The number of SVgenes found by SOMDE in different datasets.

| dataset | gene number | SV gene number | percentage |
|---|---|---|---|
| Hippocampus | 3235 | 699 | 21.61% |
| near Hippocampus | 2555 | 379 | 14.83% |
| 10X Visium brain | 14414 | 5455 | 37.85% |
| Liver | 2238 | 164 | 7.32% |
| Kidney | 4650 | 522 | 11.23% |

**Fig. S1.** Simulation results of the first set of experiments (dropout rate=0).

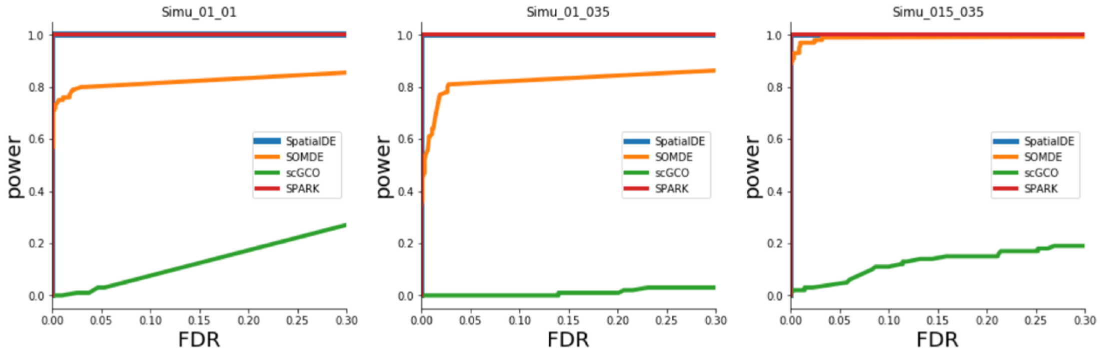

**Fig. S2.** Top50 SVgenes on 10X Visium Cancer dataset (original space and SOM space).

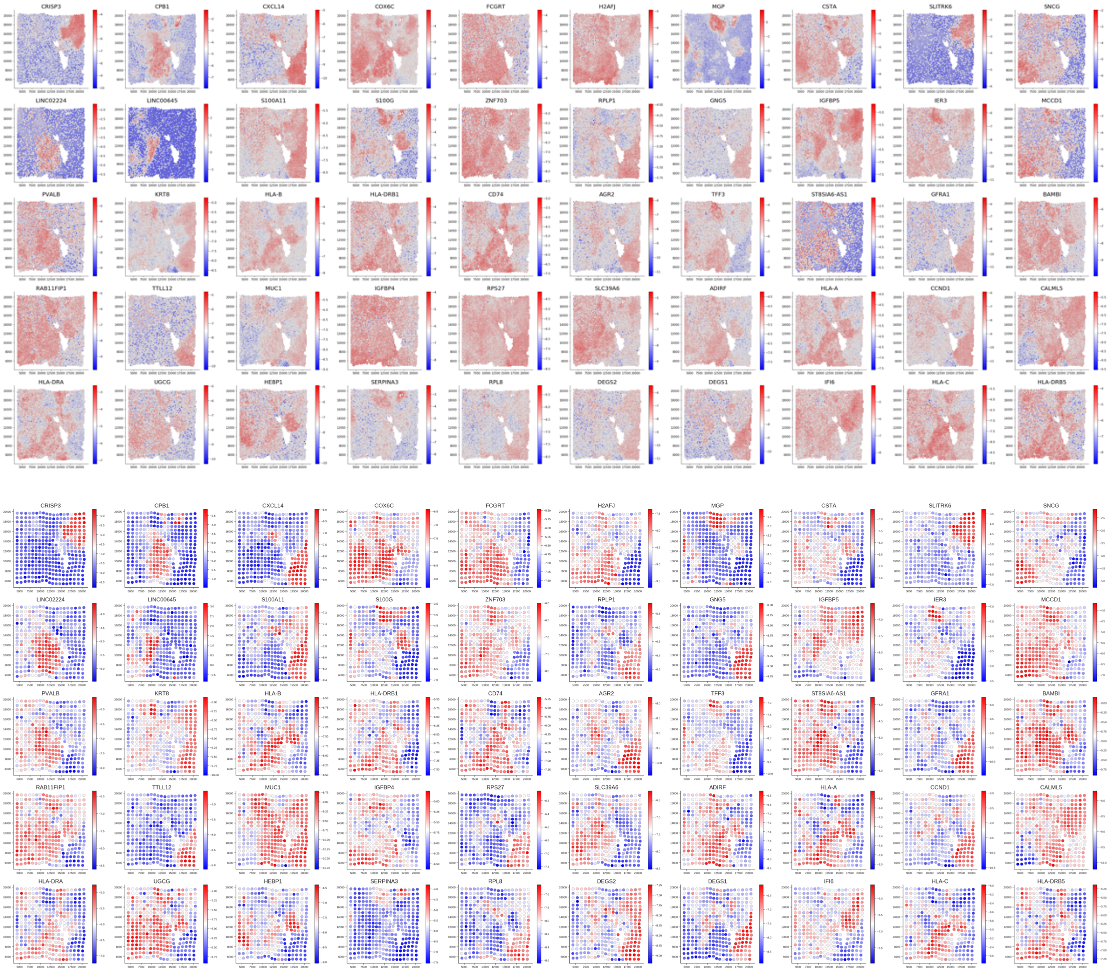

**Fig. S3.** Top50 SVgenes on 10X Visium Brain dataset (original space and SOM space).

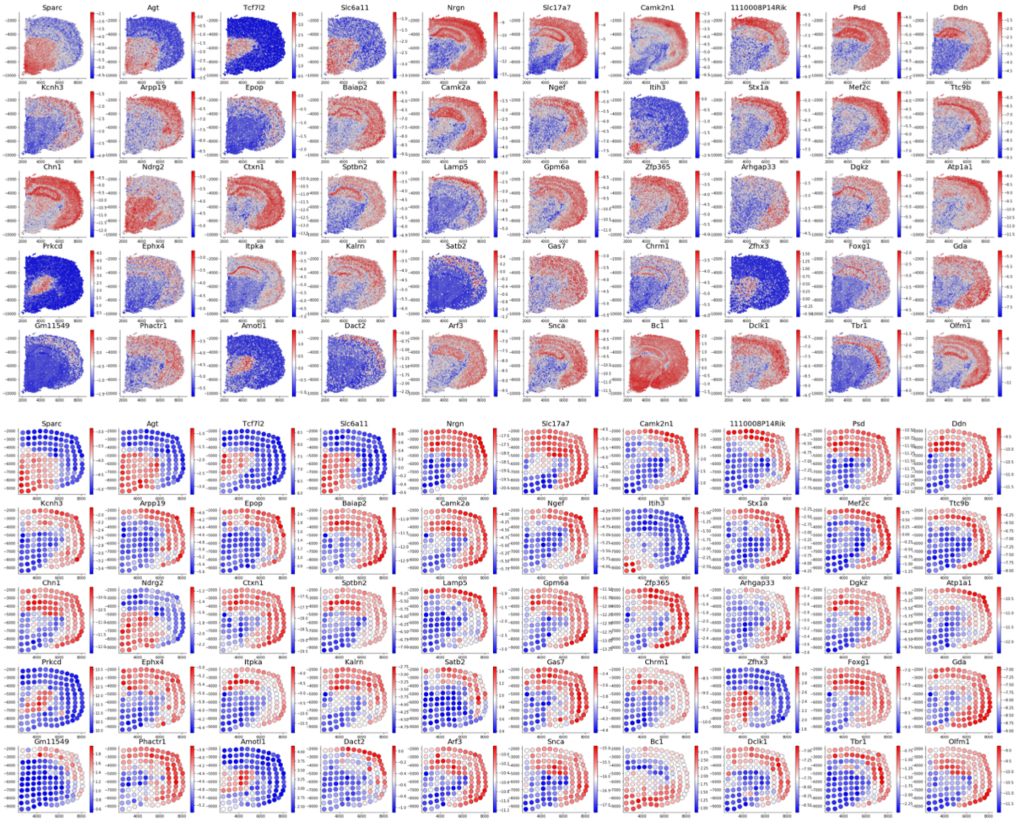

**Fig. S4.** spatial metagene found by SOMDE on 10X data. We use hierarchical clustering to obtain 9 categories from the top 1000 spatially variable genes ranked by SOMDE. The average expression of each type of gene is calculated, and these 9 genes with average expression are called metagenes. Finally, the expression is drawn on the tissue spatial domain.

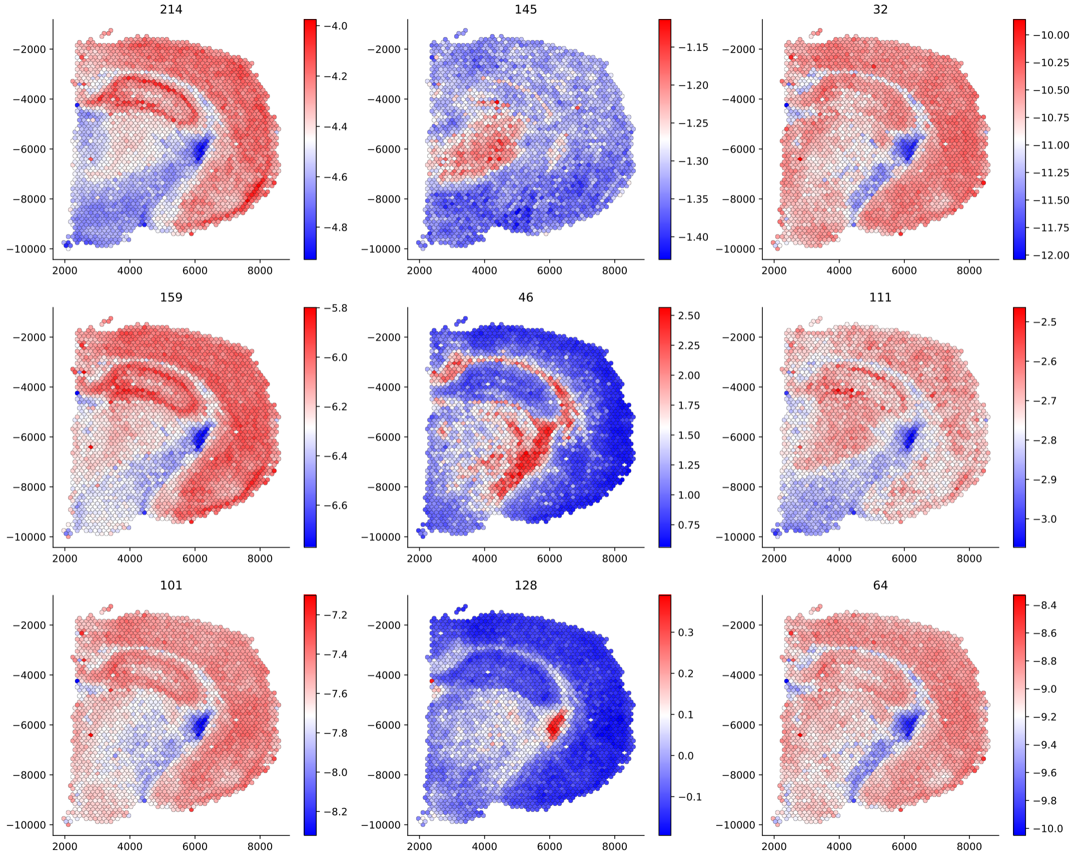

**Fig. S5.** Top50 SVgenes on Slide-seq hippocampus dataset (original space and SOM space).

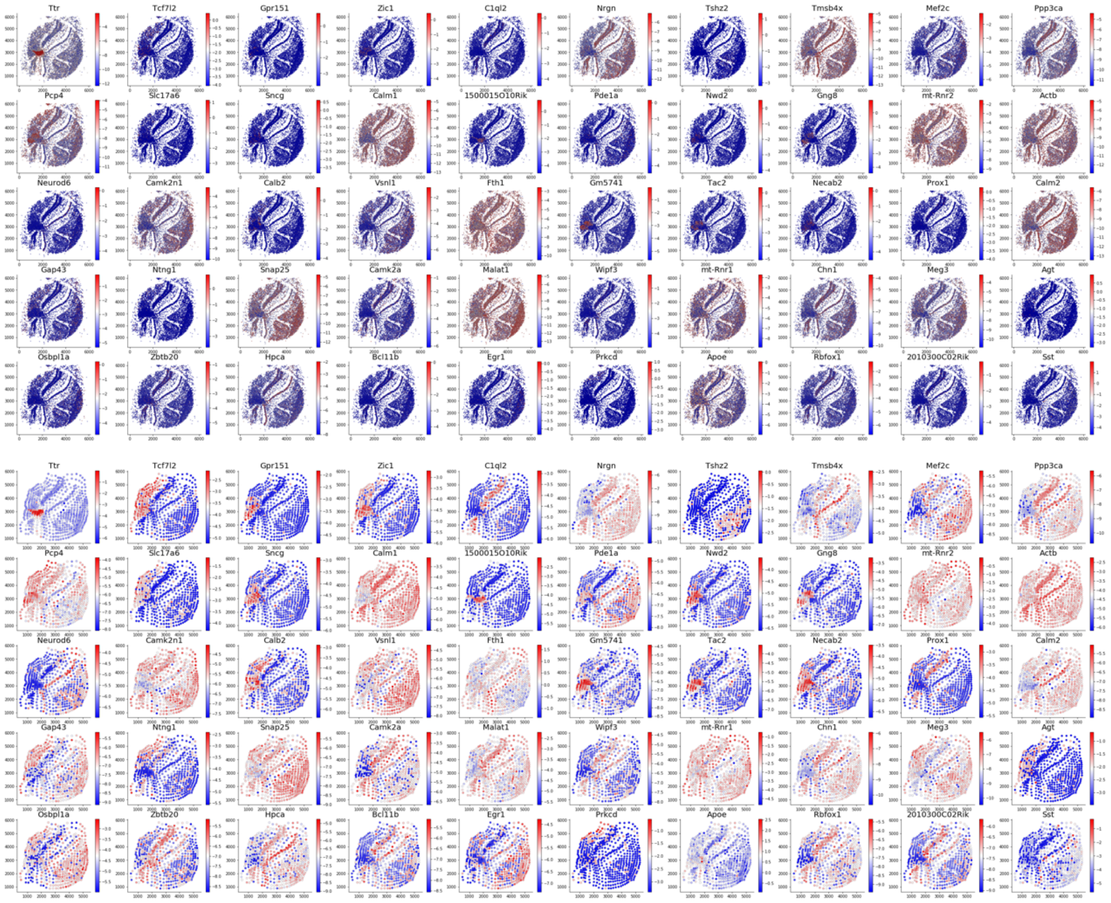

**Fig. S6.** Top50 SVgenes on Slide-seq near hippocampus dataset (original space and SOM space).

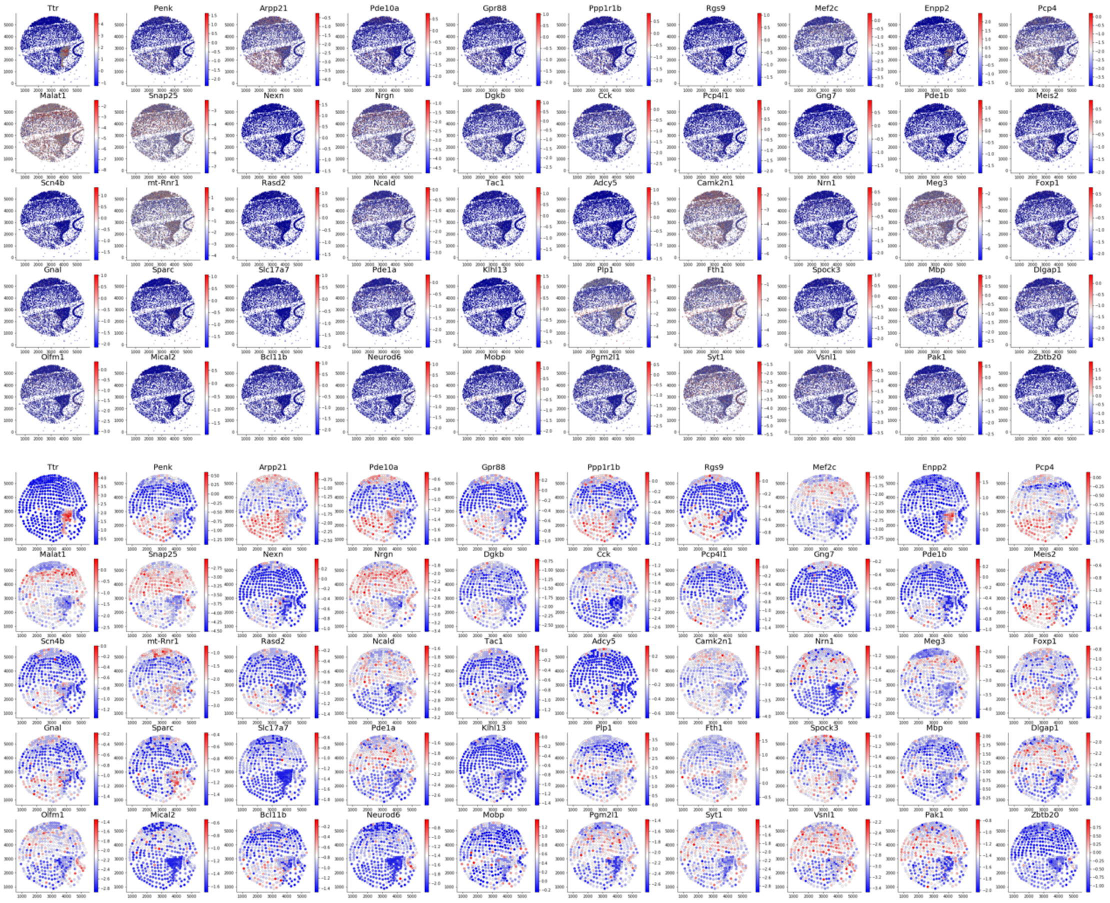

**Fig. S7.** Top50 SVgenes on Slide-seq Liver dataset (original space and SOM space).

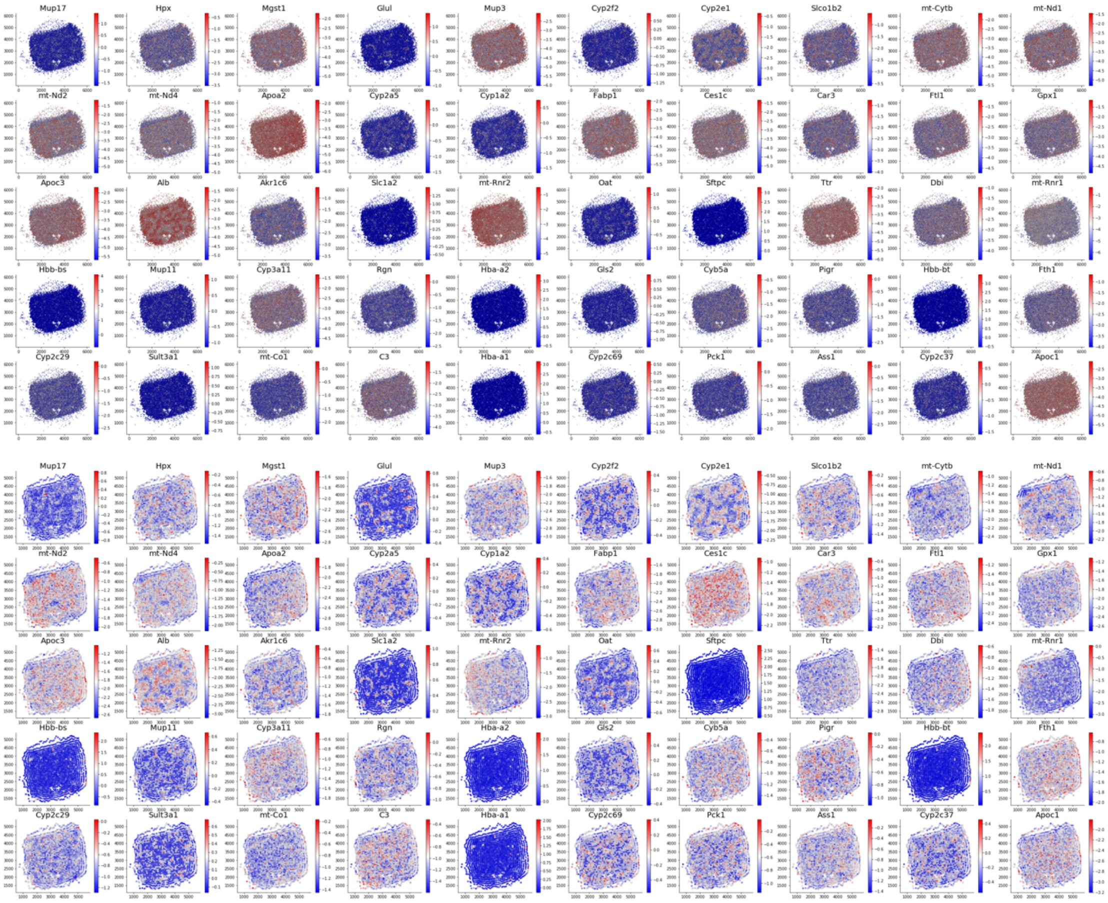

**Fig. S8.** Top50 SVgenes on Slide-seq Kidney dataset (original space and SOM space).

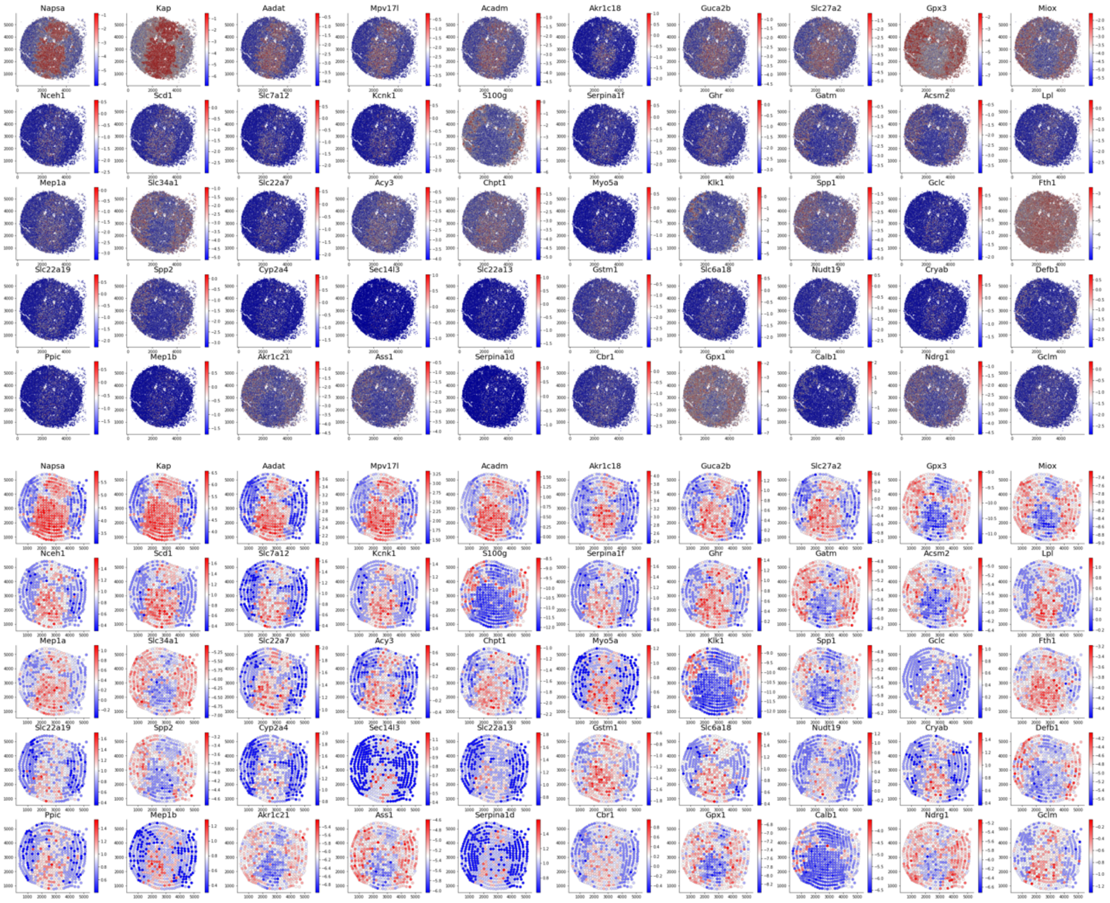

**Fig. S9.** Spatial patterns of *Nrgn* and *Camk2n1* genes and their corresponding ISH patterns.

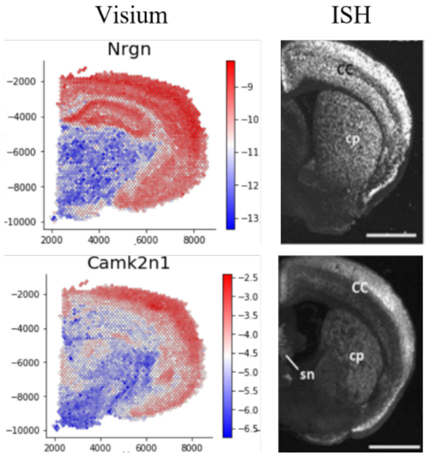


**Fig.S10.** Spatial patterns of 6 SVgenes detected in the 10X Visium data and their ISH images.

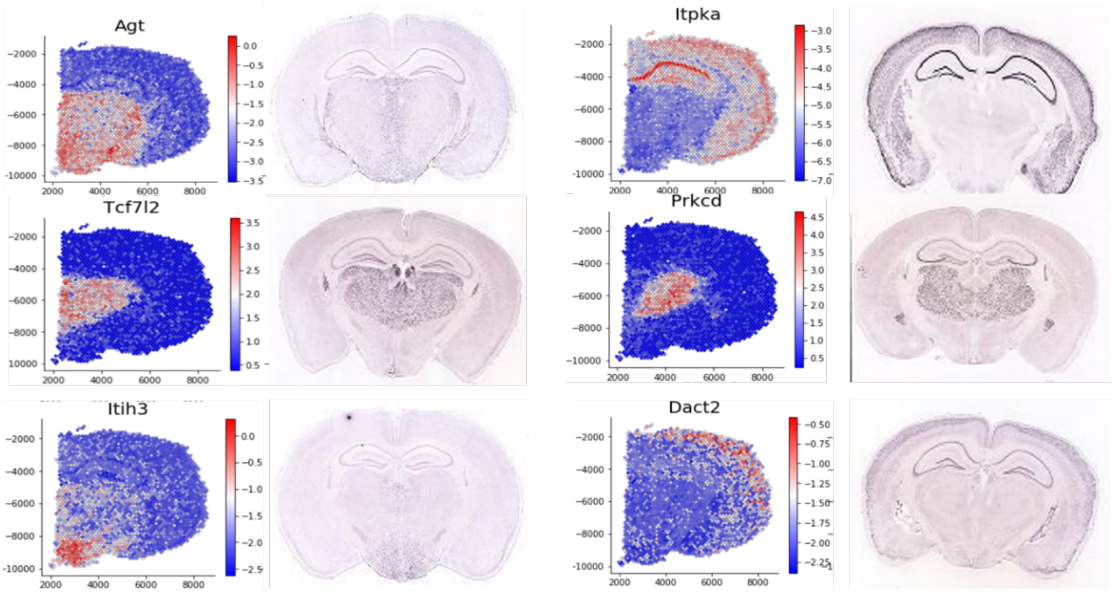

**Fig.S11.** the bottom 50 SVgenes identified by SOMDE on the 10X Brain data.

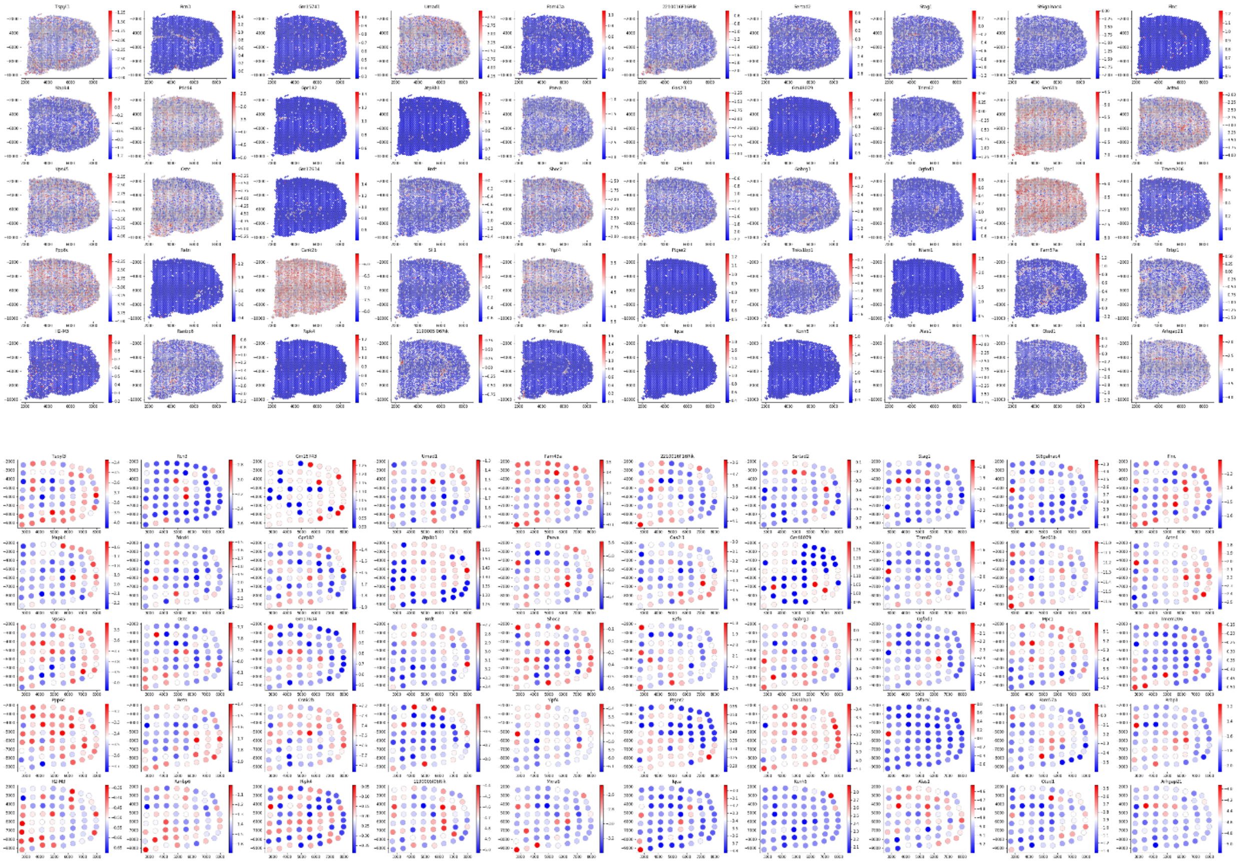

**Fig. S12.** Top 500 Genes rank similarity of SOMDE, Giotto, SPARK, cSOMDE and SpatialDE results in the 10X brain data.

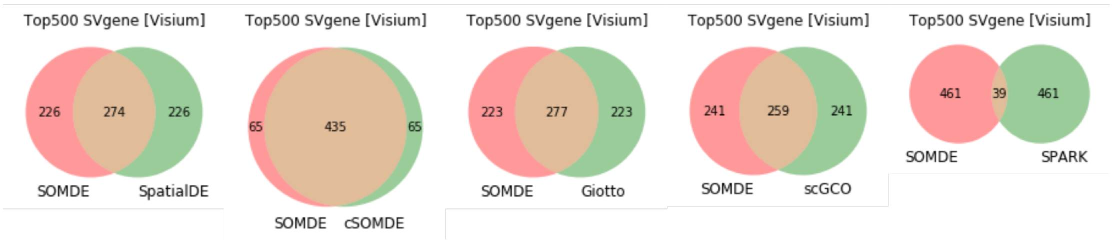


**Fig.S13.** Top 500 Genes rank similarity of SOMDE, Giotto, cSOMDE and SpatialDE results in the nHipp data.

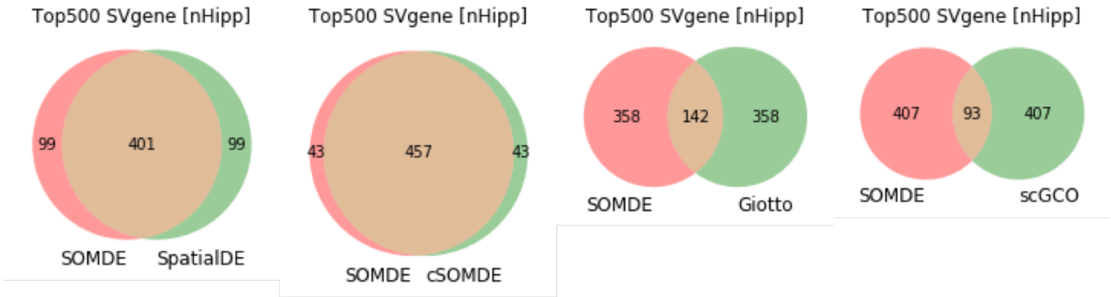

**Fig. S14.** Rank similarity of cSOMDE and SOMDE on the Visium brain and nHipp data. One spot denotes one gene, x and y coordinates are two ranks in the SOMDE and cSOMDE results, respectively.

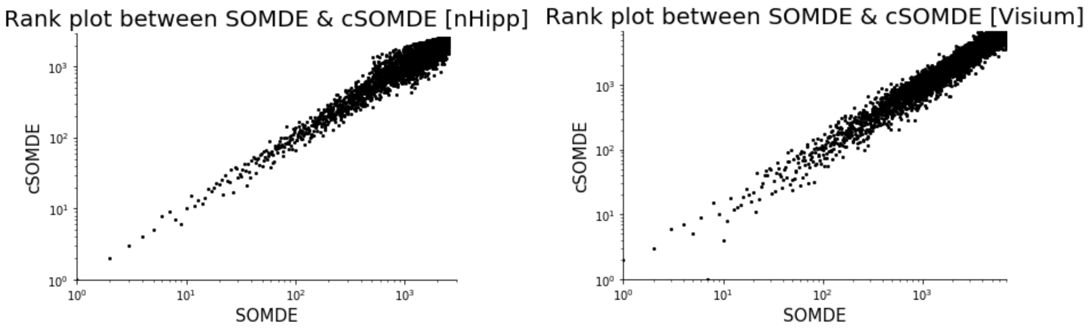


**Fig.S15.** Rank similarity of SOMDE results under different map sizes (parameter k) on the Visium brain and nHipp data.

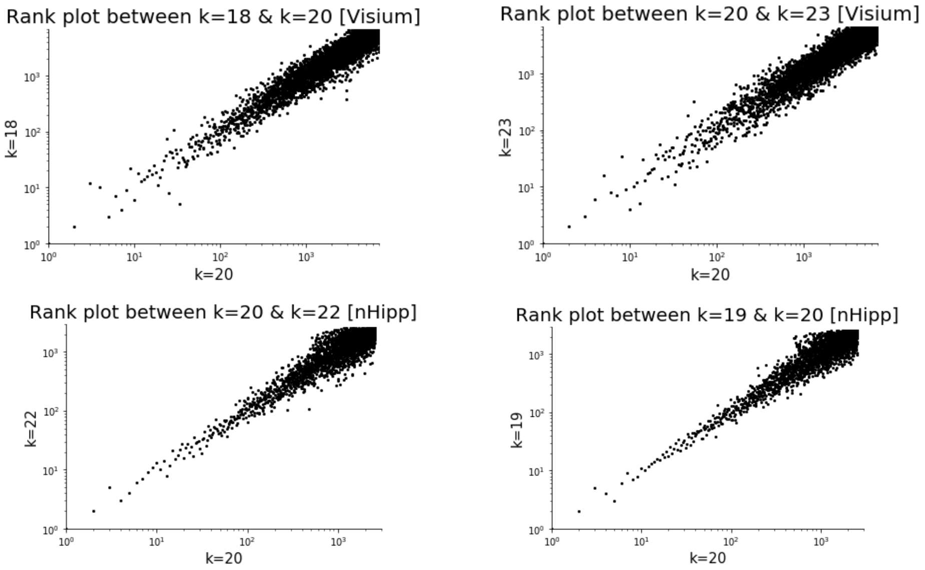

**Fig.S16.** The statistical calibration assessment of SOMDE and SpatialDE through data randomization in 10X brain data. Observed P-value are obtained from the SOMDE or SpatialDE on the randomized data, and expected P-value are sampled from the uniform distribution.

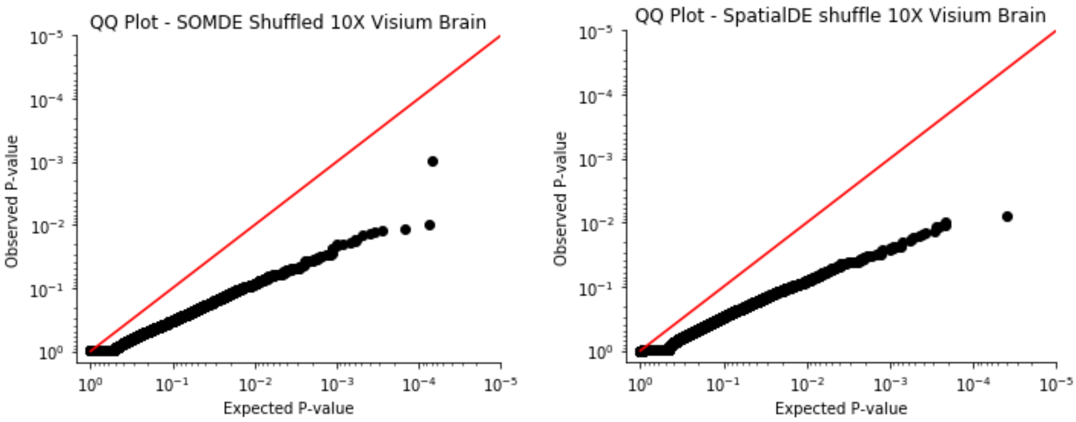


**Fig.S17**. The statistical calibration assessment of SOMDE and SpatialDE through data randomization in nHipp data.

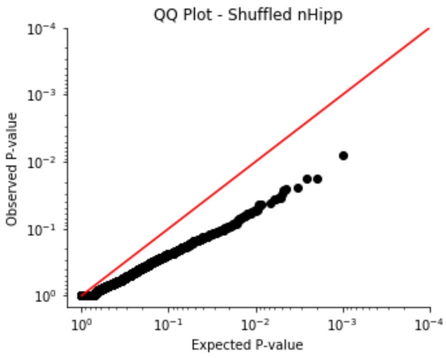


**Fig.S18**. Rank similarity of SOMDE results under different values of gamma on the nHipp data.

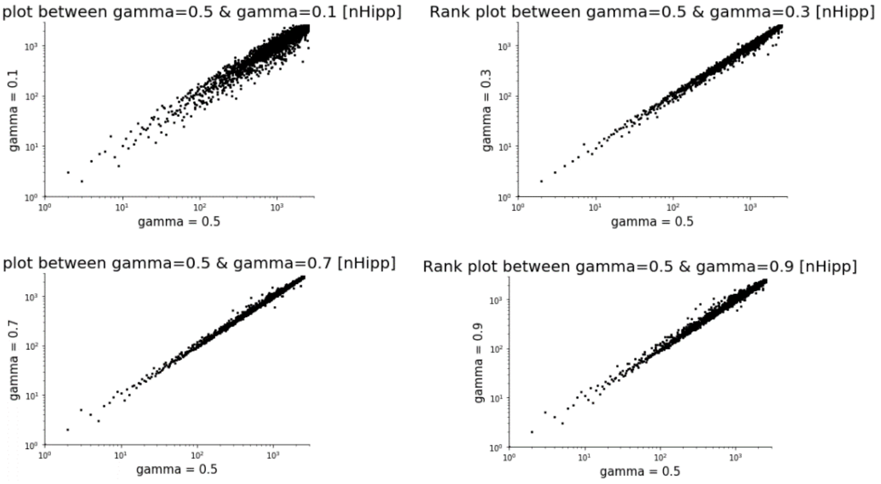

**Fig.S19.** Examples of SVgenes missed by SOMDE on the 10X brain data. They are identified by SPARK, SpatialDE, Giotto or scGCO. Genes shown in the first 3 rows are more likely to have local spatial variations but missed by SOMDE, while the gene in the last 3 rows are probably false positives but identifed by other methods. (left: expressions on the original resolution; right: expressions on the condensed maps)

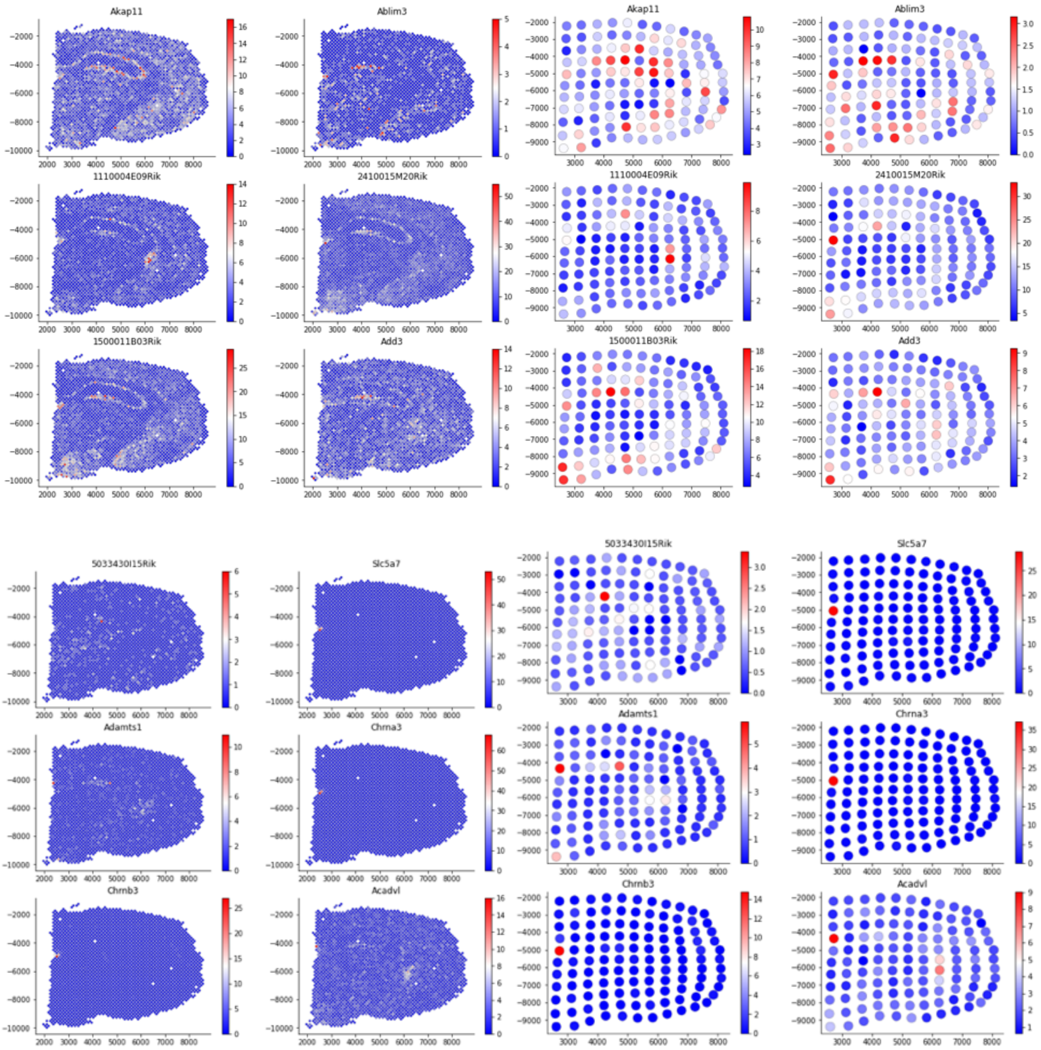
